# DNA end-resection is stimulated by an interaction between BRCA1 exon 11 and TOPBP1

**DOI:** 10.64898/2026.01.09.698382

**Authors:** Marta San Martín Alonso, Rosalie A. Kampen, Anne Schreuder, Daniel Salas, Arnoud H. de Ru, Veronica Garzero, Romy L.S. Mesman, Román González-Prieto, Peter A. van Veelen, Sylvie M. Noordermeer

## Abstract

Approximately 60% of the tumour suppressor protein BRCA1 is encoded by a single exon. Many tumours carry mutations in this exon, often resulting in exon skipping and thus a protein of a severely reduced size. Although none of the well-described protein domains of BRCA1 are encoded by this exon 11, the isoform lacking this part shows a hypomorphic activity in homologous recombination. To better understand the function of this large exon, we performed proteomic analyses to identify interaction partners via this part of the protein. Here, we report a DNA damage-and phospho-dependent interaction of TOPBP1 to the protein region of BRCA1 encoded by exon 11. Mechanistically, this interaction is required for BRCA1’s role in end-resection during homologous recombination. In contrast, the interaction is not required for TOPBP1’s role in ATR activation. Our data provide novel mechanistic insight into the function of this poorly characterized part of the BRCA1 protein.

## Introduction

Cells are constantly exposed to different exogenous and endogenous agents that can damage the DNA molecule. One of the most toxic lesions in the DNA is the DNA double-stranded break (DSB). Failure to repair such breaks can lead to genomic instability, an important cause of cancer. Fortunately, cells have evolved different ways to detect and repair damaged DNA to maintain genetic integrity [1]. The cell is equipped with multiple repair pathways to overcome DSBs, varying in their efficiency and accuracy: classical non-homologous end joining (cNHEJ), alternative end joining (aEJ), single strand annealing (SSA), and homologous recombination (HR) [2].

HR utilizes the intact sister chromatid to copy the genetic information during repair, resulting in high fidelity, error-free repair. This pathway is activated by DNA end-resection of the broken ends to expose ssDNA stretches that can be used for homology search and templated repair. BRCA1 is an important player in HR stimulating both resection and the homology search [3]. Hereditary BRCA1 mutations predispose to breast and ovarian cancers (Hall, King, Science 1990; Lord & Ashworth, Nat Rev Cancer, 2016; Couch, Fasching, JCO, 2015), but mutations also frequently arise in sporadic tumours [4]. Approximately 30% of the mutations in BRCA1 target exon 10 – historically known as exon 11 and hence also referred to as exon 11 in this manuscript - a large exon encoding for about 60% of the protein [5, 6]. Interestingly, these mutations frequently result in exon skipping (resulting in *BRCA1-Δ11*) or usage of an alternative splice donor site (*BRCA1 11q*). Both transcripts are naturally occurring isoforms coding for proteins lacking a large part of the wildtype protein although the well-studied functional domains of BRCA1 (including the RING, coiled-coil and BRCT domains) are retained, suggesting residual functions [7, 8]. Indeed, compared to BRCA1-deficiency, which is embryonically lethal, mice carrying BRCA1-Δ11 alleles are viable in a *TP53*^-/-^ background, although suffering from increased tumorigenesis [9–12]. Moreover, previous studies have demonstrated that mutations in exon 11 result in hypomorphic phenotypes such as partially reduced HR efficiency (reduced RAD51 and RPA recruitment to DNA damage sites) and resistance to therapy [13, 14]. Better understanding of the functional role of exon 11 in the maintenance of genome stability is therefore essential to improve prediction of disease progression and therapy response of patients with tumours carrying exon 11 mutations.

In this study, we identified interaction partners of the region in BRCA1 encoded by exon 11. Among these interactors was TOPBP1, which showed a damage-dependent interaction with this region. TOPBP1 is a protein consisting of eight BRCT domains and binds topoisomerase 2 β, an enzyme that relieves super-helical stress during DNA replication [15]. Additionally, TOPBP1 is able to bind DSBs and single stranded DNA (ssDNA) directly, where it can activate the ataxia telangiectasia and Rad3-related (ATR) enzyme. ATR is involved in the activation of DNA damage checkpoints, initiating cell cycle arrest to slow down the replication fork and repair the damaged DNA [16–18]. TOPBP1 is also capable to interact with NBS1, driving efficient HR [19] and is involved in balancing 53BP1 and BRCA1 interplay to stimulate HR [20]. Moreover, previous studies have identified an interaction between BRIP1 (also known as FANCJ or BACH1) and TOPBP1, specifically through the C-terminal BRCT domains (BRCT 7 and 8) of TOPBP1 [21]. BRIP1 also interacts with the BRCT domains of BRCA1 [22], and hence it has been shown that BRCA1, BRIP1 and TOPBP1 are present in the same protein complex [23]. Interestingly, our data indicate that the association between BRCA1 and TOPBP1 is only marginally affected when BRCA1 harbours a mutation that impairs its interaction with BRIP1, indicating that TOPBP1 and BRCA1 also bind in a BRIP1-independent setting. Here, we show that exon 11 of BRCA1 provides a second binding interface to TOPBP1, and that this interaction stimulates the role of BRCA1 in DNA end-resection.

## Materials and Methods

### Cell lines

All cells were kept at 37°C and 5% CO_2_ levels. BRCA1-deficient cells were kept at low O_2_ levels (3%). RPE1 hTERT and HEK 293T cells were obtained from ATCC (Manassas, VA, USA) and cultured in DMEM high glucose, Glutamax and pyruvate supplemented (Thermo Fisher Scientific, Waltham, MA, USA) supplemented with 10% FCS and 1% Penicillin/Streptomycin. RPE1 hTERT *PAC^-/-^ TP53^-/-^* and RPE1 hTERT *PAC^-/-^ TP53^-/-^ BRCA1^-/-^* (referred as *p53^-/-^* and *BRCA1^-/-^* in this manuscript) were generated by nucleofection of pLentiCRISPRv2 (Addgene: 52961 [24]) containing the following sgRNAs sg*PAC*: ACGCGCGTCGGGCTCGACAT; sgTP53: CAGAATGCAAGAAGCCCAGA; sg*BRCA1*: AAGGGTAGCTGTTAGAAGGC. Subsequently, cells were clonally expanded and genotyping was performed by PCR amplification and Sanger sequencing of the targeted locus, followed by TIDE analysis [25]. RPE1 hTERT *TP53^-/-^ BRCA1^-/-^* and the isogenic *TP53^-/-^ BRCA1^-/-^ 53BP1^-/-^* cell lines used in Figure S3e have been described before [26].

The BRCA1 gene in RPE1 hTERT *PAC^-/-^ TP53^-/-^* cells was endogenously C-terminally tagged with a triple TY-1 tag using electroporation of the pLentiCRISPR-v2 vector with a BRCA1 sgRNA: 5’-TGGCTGGCTGCAGTCAGTAG-3’ and the C-terminal tagging vector BRCA1-XhoI-tag-P2A-Neo with 500 bps homology arms annealing to BRCA1’s C-terminus, a triple TY-1-tag, P2A and a neomycin resistance cassette derived from the previously described YFP-P2A-Neo vector [27]. Subsequently, cells were clonally expanded and genotyping was performed by PCR amplification and Sanger sequencing of the targeted locus.

SUM149PT cells (kind gift from Prof. D. Durocher, Lunenfeld Tanenbaum Research Institute) were cultured in Ham’s F-12 nutrient mix (Thermo Fisher Scientific) supplemented with 5 % FCS, 1% of 1M HEPES, 5 µg/mL insulin (I9278-5ML, Sigma-Aldrich, Burlington, MA, USA), and 1 µg/mL hydrocortisone (H4001-1G, Sigma-Aldrich).

A mouse embryonic stem cell line (mES) was used to express the full-length human BRCA1 gene and two gene variants with deletions comprising the coding region of exon 11 (ΔE11 (r.671_4096del) and ΔE11q (r.788_4096del) located on a Bacterial Artificial Chromosome (BAC RP11-812O5). The *BRCA1* gene variants were generated using a positive/negative selection procedure in combination with Red/ET BAC recombination as described previously [28]. The three BACs containing either the WT gene or exon 11 deletions were transfected in a hemizygous mouse Brca1 mES line (Brca1 -/loxP Pim1DR-GFP/WT 129/Ola E14 IB10) [29] using 6 μg of BAC DNA to transfect 5 x 10^6^ cells per variant using Lipofectamine 2000 (Invitrogen, Thermo Fisher Scientific). Transfected cells were seeded in 10-cm cell culture dishes and cultured in the presence of G418 (200 μg/ml), starting 24 h post transfection. Thirteen days post transfection cells were treated for 16 hours with 1.0 µM 4-Hydroxytamoxifen (4-OHT) to remove the mBrca1 conditional allele. After culturing the mES cells for six days in BRL-conditioned mES cell culture medium containing puromycin (1.8 µg/mL), experiments was conducted. Mouse mES cells were cultured in DMEM/F-12 medium (Thermo Fisher Scientific) supplemented with 10% FCS, 1% Penicillin–Streptomycin (5000 U/mL, Thermo Fisher Scientific), 5 mg/mL insulin (Sigma-Aldrich), 5 ng/mL epidermal growth factor (Thermo Fisher Scientific, 53003-018), and 5 ng/mL cholera toxin (Sigma-Aldrich, C8052).

All cells were regularly checked for mycoplasma contamination.

### Antibodies

The following primary antibodies were used for western blotting: Mouse α BRCA1 (Merck, OP92; 1:1000), Rabbit α BRCA1 (Merck, 07-434; 1:1000), Rabbit α BRIP1 (Sigma-Aldrich, B1310-200; 1:2000), Mouse α TUBULIN (Sigma-Aldrich, T6199; 1:5000), Rabbit α TOPBP1 (Abcam, ab2402; 1:1000), Rabbit α p-CHK1 (Cell signalling Technology, 2348S; 1:1000), Rabbit α CHK1 (Abcam, ab47574; 1:1000), Mouse α TY-1 (Diagenode, C1520054; 1:1000), and Rabbit α GAPDH (Sigma-Aldrich, G9545; 1:5000), Rabbit α p-ATR (Abcam, ab223258, 1:1000), Rabbit α ATR (Bethyl laboratories, A300-138A, 1:5000), Rabbit α p-RPA-S33 (Bethyl laboratories, A300-246A, 1:1000), Mouse α RPA (Abcam, ab2175, 1:1000), Mouse α γH2AX (Millipore, 05-636, 1:1000).

The following secondary antibodies were used for western blotting: Goat α Mouse or Goat α Rabbit labelled with respectively IRDye 800 or IRDye 680 (LI-COR; 1:15000) and HRP-labelled Goat α Mouse and Donkey α Rabbit (Thermo Fisher Scientific; 1:5000).

The following primary antibodies were used for immunofluorescence: Rabbit α BRCA1 (Millipore, 07-434; 1:1,000), Mouse α BrdU (Amersham, RPN202; 1:800), Rabbit α TOPBP1 (Abcam, ab2402; 1:1000), Rabbit α p-RPA32 (Ser4/Ser8) (Bethyl laboratories, A300-245A; 1:1000), Mouse α TY-1 (Diagenode, C1520054; 1:1000), and Rabbit α RAD51 (Bio Academia, 70-001; 1:15000). The following secondary antibodies were used for immunofluorescence: Goat α Mouse or Goat α Rabbit labelled with either Alexa 488 or Alexa 555 (Thermo Fisher Scientific, Invitrogen; 1:1000).

### Plasmids & cloning

Purification of plasmid DNA or PCR products was performed using commercially available kits (Qiagen, Hilden, Germany) according to the manufacturer’s protocol. sgRNAs were cloned into BsmBI-digested pLentiCRISPRv2. To obtain pCW57.1-BRCA1-3xTY-1, full length BRCA1 was PCR-amplified from pCL-MFG-BRCA1 (Addgene, 12341; (Ruffner and Verma, 1997)) with a triple TY-1-tag included in the reverse primer and subsequently cloned into pENTR_1A (Thermo Fisher Scientific) and transferred to pCW57.1 (Addgene, 41393) via gateway cloning. To obtain point mutations in BRCA1, we used site directed mutagenesis in the plasmid pENTR_1A_BRCA1-3xTY-1 followed by gateway reaction to introduce the BRCA1 variant into pCW57.1. To get deletions in BRCA1, PCR and Gibson assembly (E5510S, NEB) was performed in the plasmid pENTR_1A_BRCA1-3xTY-1 followed by gateway cloning into pCW57.1.

To obtain a pCW57.1-FLAG-TOPBP1 plasmid, TOPBP1 CDS was obtained via PCR from the plasmid pcDNA3-LacR-TopBP1 (Addgene, 31317; [30]) and cloned into the plasmid pDONR221. The resultant plasmid was subjected to site directed PCR mutagenesis to create silent mutations at the genomic locus of the gRNA used for the depletion. Finally, a gateway reaction was performed to introduce the TOPBP1 cassette into pCW57.1-Zeo. pCW57.1-Zeo was created by exchanging the puro resistance cassette with a Zeocin cassette into the plasmid pCW57.1 (using restriction digestion with AscI and NheI). To create TOPBP1 deletion variants, PCR and subsequent Gibson assembly was performed in the plasmid pCW57.1-FLAG-TOPBP1.

### Viral transductions and transfections

Lentivirus were produced in HEK 293T cells by jetPEI transfection (Polyplus Transfection, Illkirch, France) of pLentiCRISPRv2 or pCW57.1 plasmids with third generation packaging vectors pMDLg/pRRE, pRSV-Rev and pMD2.G. Viral supernatants were harvested 48-72 hours post transfection and used to transduce cells at an MOI of ∼ 1 in the presence of 4 µg/mL polybrene. For RPE1 cells with an intact PAC gene, 10 µg/mL puromycin was used for selection 48 hr post-transduction, for RPE1 *PAC^-/-^* cells, 1-2 µg/mL of puromycin was used. For zeocin selection, 200 µg/mL were used. Transfections with siRNAs were performed using Lipofectamine RNAiMAX (Invitrogen, Carlsbad, CA, USA) according to the manufacturer’s protocol. Transfected cells were used for experiments 48-72 hours after transfection.

### Western Blotting

Cells were lysed in RIPA lysis buffer (1% NP-40, 50 mM Tris-HCl pH 7.5, 150 mM NaCl, 0.1% SDS, 3 mM MgCl_2_, 0.5% sodium deoxycholate) supplemented with cOmplete Protease Inhibitor Cocktail EDTA-free (Sigma-Aldrich) and 100 U/mL Benzonase (Sigma-Aldrich). 4 x BOLT LDS sample buffer (Thermo Fisher Scientific) with 10x BOLT sample reducing agent containing DTT (Thermo Fisher Scientific) was added to the lysates to obtain 1 x dilutions, followed by denaturation at 95°C for 5 minutes. Proteins were separated by SDS-PAGE on 4-12 % gradient gels (Thermo Fisher Scientific) and transferred to Amersham Protran premium 0.45 µm nitrocellulose membrane (GE Healtcare Life Sciences, Chicago, IL, USA). Membranes were blocked with 8% skimmed milk (Santa Cruz Biotechnology, Dallas, TX, USA) or with Blocking buffer for fluorescent western blot (Rockland, Pottstown, PA, USA), and stained with primary and secondary antibodies in the same buffer used for blocking. After secondary antibody staining, membranes were imaged on an Odyssey CLx scanner (LI-COR BioSciences, Milton, UK), followed by image analysis using ImageStudio (LI-COR BioSciences). Alternatively, when HRP-labelled secondary antibodies were used, the membranes were treated with the WesternBright ECL HRP Substrate kit (Advansta, San Jose, CA, USA) and imaged on an Amersham Imager 680 (Bioké, Leiden, The Netherlands) or iBright (Thermo Fisher Scientific).

### Immunoprecipitation

Cells were grown in 2 or 3 15-cm dishes up to 90% confluency. 24 hr after seeding cells were treated with dox for 48 hr. Cells were collected 1 hr post IR (5Gy) in 1mL ice-cold NETT buffer (100 mM NaCl, 50 mM TRIS pH 7.6, 5 mM EDTA pH 8.0) supplemented with 1x cOmplete protease inhibitor cocktail EDTA-free (Sigma-aldrich), 0.5% Triton X-100, 7 mM MgCl_2_ and 500 U of benzonase (EMD millipore). After cell lysis and clearance by centrifugation (14000 rpm, 10 min), supernatants were incubated for 4 hours with 20 μL Dynabeads protein-G (Thermo Fisher Scientific) that were pre-incubated for 2 hours and 30 minutes with 2.2 μg α-TY-1 antibody (Diagenode; C1520054) to immunoprecipate BRCA1. To immunoprecipitate TOPBP1, lysates were incubated with 2 μg α-TOPBP1 (Abcam; ab2402) and 20 μL Dynabeads protein-A (Thermo Fisher Scientific). For FLAG-IP, lysates were incubated with 2 μg α-FLAG (Sigma-Aldrich; F1804) and 20 μL Dynabeads protein-G.

In the case of subsequent SDS-PAGE and western blotting, lysates were washes 5 times with NETT buffer and denatured with 2x Bolt LDS sample buffer + reducing agent and incubated at 95°C for 5 minutes before loading.

### LC-MS/MS

#### On-bead digestion

After TY-1 immunoprecipitation, samples were washed 3 times with NETT buffer and 4 times with 50 mM ammonium bicarbonate (ABC) prepared in HPLC grade (keratin free) water. For reduction and alkylation of the samples, 10 mM DTT (prepared in 50 mM ABC) was added to the beads and incubated 25 minutes at 60°C. Next, 15 mM 2-Chloroacetamide (2CAA) (prepared in 50 mM ABC) was added to the beads and incubated 25 minutes at room temperature in the dark. Afterwards, 0.8 µl of 500 mM DTT was added to neutralise the 2CAA and incubated 5 minutes at room temperature.

For the trypsin digestion, 160 µl 50 mM ABC and 1 μL Trypsin (1mg/mL stock) was added to the beads and incubated overnight at 37°C while nutating. Afterwards, 0.5 μL Trypsin was added and incubated for an additional 3 hours at 37°C while nutating. Beads were washed 2 times with 100 μL ABC, and supernatants were pooled to a new tube, followed by a centrifugation at 10000 g. Samples were dried in a speed vacuum and stored at −20 °C.

#### Stage Tip clean-up

Stagetips (C18, SP-301, Thermo Fisher Scientific) were prewetted with 50 μl 100% Methanol and centrifuged 2 minutes at 700 g. 50 μL of buffer B (80% Acetonitrile, 1% TFA, 0.5% Acetic Acid) was added to each stagetip and centrifuged 2 minutes at 700 g. 50 μl of buffer A (1 % TFA, 0.5 % Acetic Acid in HPLC (keratin-free) water) was added and centrifuged 2 minutes at 700 g. After preparation of the stagetips, freeze-dried peptides were dissolved in 50 μl 1 % TFA and centrifuged at maximum speed for 10 minutes. Afterwards, dissolved peptides were added to the equilibrated stagetips and passed by low-speed centrifugation (5 minutes at 100 g). Next, stagetips were washed with 50 μl of buffer A and centrifuged at 150 g per 5 minutes. Peptides were eluted using 20 μl of buffer B and centrifuged 2 minutes at 100 g. These last two steps were repeated and next, peptides were speed vacuumed until complete dryness and stored for Mass spectometry.

#### Mass Spectrometry

Peptides were dissolved in 0.1% formic acid and subsequently analyzed by online C18 nano-HPLC MS/MS with a system consisting of an Easy nLC 1200 gradient HPLC system (Thermo Fisher Scientific), and an Orbitrap Fusion LUMOS mass spectrometer (Thermo Fisher Scientific). Fractions were injected onto a homemade precolumn (100 μm × 15 mm; Reprosil-Pur C18-AQ 3 μm, Dr Maisch, Ammerbuch, Germany) and eluted via a homemade analytical nano-HPLC column (30 cm × 75 μm; Reprosil-Pur C18-AQ 1.9 μm). The analytical column temperature was maintained at 50°C with a PRSO-V2 column oven (Sonation, Biberach, Germany). The gradient was run from 2% to 36% solvent B (20/80/0.1 water/acetonitrile/formic acid (FA) v/v) in 120 min. Solvent A was water/formic acid 100/0.1 (v/v). The nano-HPLC column was drawn to a tip of ∼5 μm and acted as the electrospray needle of the MS source.

The mass spectrometer was operated in data-dependent MS/MS mode with a cycle time of 3 seconds, with the HCD collision energy at 32% and recording of the MS2 spectrum in the orbitrap, with a quadrupole isolation width of 1.2 Da. In the master scan (MS1) the resolution was 120,000, the scan range 400-1500, at standard AGC target at maximum fill time of 50 ms. A lock mass correction on the background ion m/z=445.12003 was used. Precursors were dynamically excluded after n=1 with an exclusion duration of 45 s, and with a precursor range of 20 ppm. Charge states 2-5 were included. For MS2 the first mass was set to 110 Da, and the MS2 scan resolution was 30,000 at an AGC target “standard” at maximum fill time of 60 ms.

### Mass Spectrometry data analysis

All raw data were analyzed using MaxQuant (version 2.1.3.0) as previously described [31]. We performed the search against an in silico digested UniProt reference proteome for Homo sapiens including canonical and isoform sequences (24th January 2022). Database searches were performed according to standard settings after reduction and alkylation of samples: Oxidation (M) and acetyl (Protein N-term) were allowed as variable modifications while Carbamidomethyl (C) was allowed as fixed modification, keeping the maximum number of modifications per peptide equal to 3. Digestion was performed with Trypsin/P, allowing a maximum of 4 missed cleavages. Label-Free Quantification (LFQ) was enabled, not allowing Fast LFQ while permitting iBAQ and matching between runs.

Output from MaxQuant Data was exported and processed for statistical analysis in the Perseus computational platform version 2.0.7.0 [32]. LFQ intensity values were log2 transformed and potential contaminants and proteins either identified by site or only reverse peptides were removed. Samples were grouped in experimental categories and the matrix was separated in two for further filtering. For the exogenous BRCA1-3xTY-1 samples, proteins not identified in 3 replicates in at least one group were removed. For the endogenous BRCA1-TY-1 IP runs, proteins not identified in 2 out of 3 replicates in at least one group were removed. Missing values were imputed using normally distributed values with 0.3 width and 1.8 down shift for the total matrix. After imputation, Student’s t tests with a threshold p-value of 0.05 and S0=1 for both sides were applied. Statistical analysis tables were exported and processed in MS Excel. Volcano plots were constructed using VolcaNoseR [33] web app with a significant threshold of 1.3 for exogenous BRCA1-TY-1 and 0.7 for endogenous BRCA1-TY-1 according to a p-value of 0.05 and 0.2 respectively. A supplemental text file is available describing the Maxquant/Perseus output data provided as supplementary tables.

### Immunofluorescence

Cells were grown with dox in 24 well plates containing 13 mm glass coverslips up to 85% confluency, treated with Dox for 48 hr if required, and irradiated with 10 Gy to study IR-induced foci. For p-RPA and BrdU foci analyses, cells were pre-extracted after 4 hours post-IR with cold nuclear extraction (NuEx) buffer (20 mM HEPES pH 7.5, 20 mM NaCl, 5 mM MgCl_2_ supplemented with 1mM DTT, 0.5% NP-40 (Sigma-Aldrich), 1x cOmplete protease inhibitor cocktail (Sigma-Aldrich)) for 12 minutes at 4°C and fixed with 4% paraformaldehyde in PBS for 20 minutes at room temperature and gentle shaking.

For RAD51 and BRCA1 foci observation, 4 hours after irradiation, cells were fixed and permeabilized with 1% paraformaldehyde, 0.3% Triton-X100 in PBS for 20 minutes at room temperature, follow by a second fixation step with 1% paraformaldehyde, 0.3% Triton-X100 and 0.5% methanol in PBS for 20 minutes at room temperature.

After fixation, cells were washed three times with PBS and blocked with PBS^+^ (5g/L BSA, 1.5 g/L glycine in PBS 1x). Antibody dilutions were made in PBS^+^ and incubated at room temperature. After 5 washes with PBS, cover slips were mounted using Aqua Poly/Mount (Polysciences, Warrington, USA). Images were taken using Zeiss Axio Imager 2 fluorescent microscope with a 40x zoom. Foci of at least 100 cells per condition per replicate were quantified using the IRIF analysis 3.2 Plugin in ImageJ [34].

### Clonogenical survival assay

RPE1 hTERT cells were seeded in 10-cm dishes (RPE1 WT or RPE1 BRCA1^-/-^ complemented with BRCA1-WT: 250 cells; RPE1 BRCA1^-/-^ or complemented with EV: 1500 cells, RPE1 BRCA1^-/-^complemented with BRCA1-Δ11: 500 cells) and treated as indicated. Medium containing olaparib (16 nM) (Selleck Chemicals, Planegg, Germany) and dox (1 µg/mL) was refreshed after 7 days. After 14 days, colonies were stained with crystal violet (0.4 % (w/v) crystal violet, 20% methanol) and counted manually.

### Short term proliferation assays

Cells were seeded into 6-well format in the presence of dox and olaparib (1000 nM) in complete culture medium (DMEM + 10% FCS + Pen/Strep) in duplicate. Cells were transfected with siControl (Thermo Fisher Scientific, Invitrogen; 4390843) or siTOPBP1 (Thermo Fisher Scientific, Invitrogen; Assay ID 108170) and after 7 days, cells were washed with PBS, trypsinised and collected. Viable cells were counted using a Vi-CELL XR Cell Viability Analyzer (Beckman Coulter).

## Results

### BRCA1 binds TOPBP1 through a region encoded by exon 11

To better understand the function of exon 11 of BRCA1 during DSB repair, we aimed to identify proteins that bind BRCA1 specifically through the protein region encoded by this exon 11. For this, we generated an isogenic cell line model expressing either full length (FL) BRCA1 or a variant lacking exon 11 (Δ11) by virally reconstituting RPE1 hTERT *TP53^-/-^ BRCA1^-/-^* cells with doxycycline (dox) inducible versions of the two BRCA1 protein variants with C-terminal triple TY-1 tags (or an empty vector control) (Figure 1a). Compared to FL BRCA1, BRCA1-Δ11 was expressed at higher levels, as has been observed before in other cell line models and clinical samples [13, 35]. Phenotypic characterization of these cell lines confirmed the hypomorphic effect of BRCA1-Δ11 [13, 14], showing reduced DNA end-resection (Figure 1b and S1a), reduced RAD51 foci formation (Figure 1c and S1b), mild PARPi sensitivity (Figure 1d), and no significant difference in DNA damage recruitment compared to the FL version of BRCA1 (Figure 1e).

**Figure 1.**
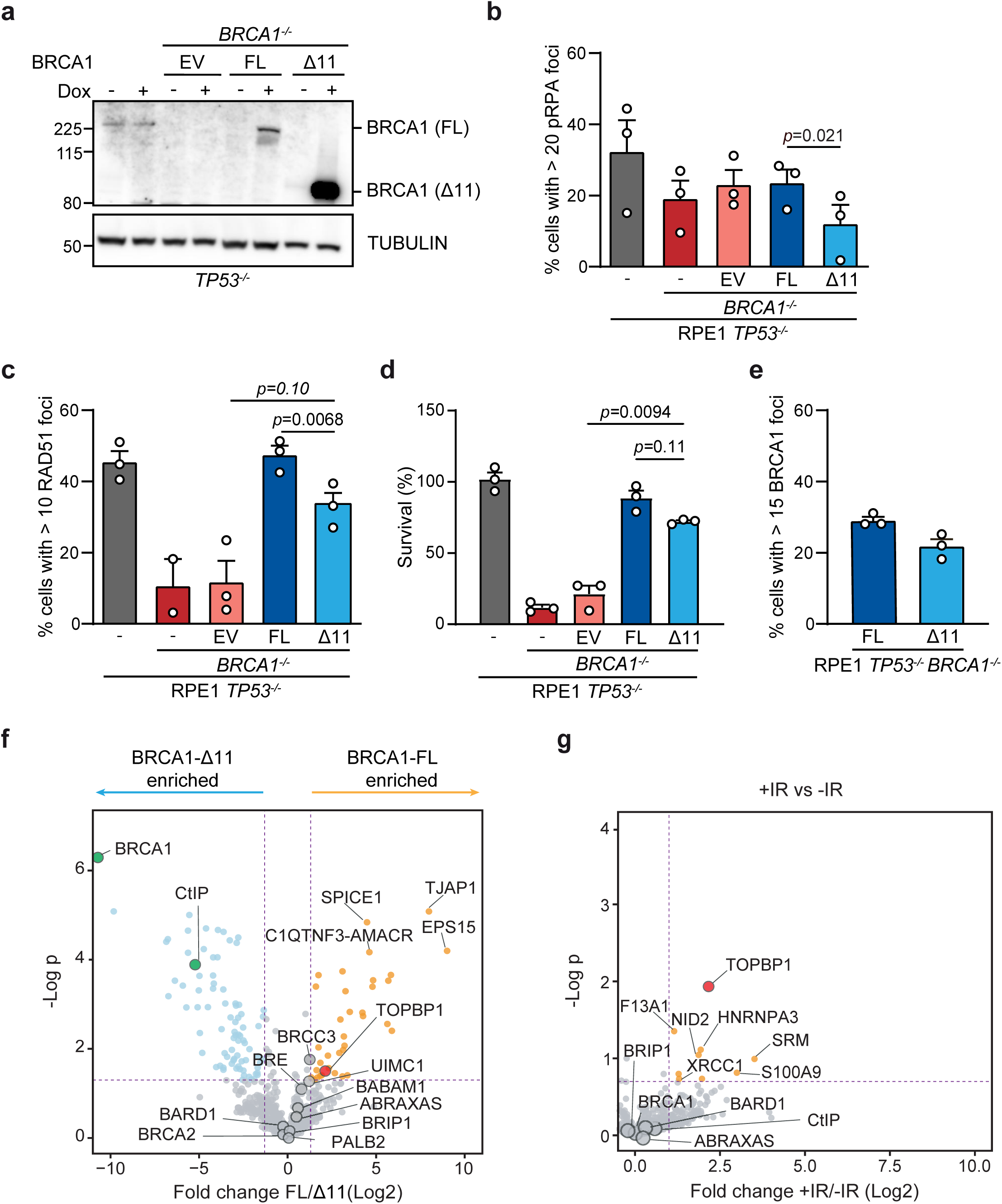
BRCA1 binds TOPBP1 through its exon 11 encoded region. (A) Lysates of RPE1 hTERT cell lines either *TP53^-/-^* or *TP53^-/-^ BRCA1^-/-^* virally reconstituted with a dox-inducible BRCA1-TY-1 cDNA (FL), BRCA1-TY-1 exon11-less (Δ11) cDNA or an empty vector (EV) control analyzed by western blotting to indicate BRCA1 variant expression 24 hr post dox induction (1 μg/mL). (B) *TP53^-/-^* or *TP53^-/-^ BRCA1^-/-^* virally reconstituted with dox-inducible FL BRCA1-TY-1 or BRCA1-TY-1-Δ11 were irradiated with 10 Gy and 4 hr post-IR analyzed by IF microscopy to detect p-RPA foci formation (n=3, mean + SEM, paired t-test, p-values are shown). A minimum of 100 cells were counted per replicate. (C) Same cells used in (A) and (B) were irradiated as in (B) and IF microscopy was performed to quantify RAD51 foci (n=2/3, mean + SEM, paired t-test, p-values are shown). A minimum of 100 cells were counted per replicate. (D) Clonogenical survival assay with continuous treatment of 16 nM olaparib and 1 μg/mL dox of the same cell lines described in (A). Data is normalized to cells treated with dox, but not treated with PARPi (n=3, mean + SEM, paired t-test, p-values are shown). (E) Same cells used in (A) and (B) were irradiated as in (B) and IF microscopy was performed to quantify BRCA1 foci (n=3, mean + SEM). A minimum of 100 cells were counted per replicate. (F) Volcano plot depicting statistical differences in binding proteins identified by MS proteomic analysis upon TY-1 IP of RPE1 hTERT *TP53^-/-^ BRCA1^-/-^* cells virally reconstituted with dox-inducible BRCA1-FL or BRCA1-Δ11 1 hr post 5 Gy IR (n=3). (G) Volcano plot depicting statistical differences in binding proteins identified by MS proteomic analysis upon TY-1 IP of mock- or IR-treated (1 hr post 5 Gy) RPE1 hTERT *TP53^-/-^* cells with an endogenous BRCA1-TY-1 tag.

To identify interaction partners of exon 11’s coding region, we performed immunoprecipitation (IP) experiments followed by liquid chromatography-tandem mass spectrometry (LC-MS/MS) (IP-MS) analyses. After irradiating the cells to induce DSBs, we purified FL BRCA1 or BRCA1-Δ11 and their interactors using antibodies against the TY-1 tag. Using this approach, we identified many well-described interactors that bind BRCA1 via domains outside of exon 11 (e.g. BARD1, PALB2, ABRAXAS, BRCA2, CtIP, BRIP1). Indeed, all these interactors were retained in BRCA1-Δ11 (Figure 1f, S1c and Supplemental Text and Supplemental Table 1). Importantly, we observed a set of proteins in our FL BRCA1 IP which were not identified in the BRCA1-Δ11 IP. Given the low expression level of FL BRCA1 compared to BRCA1-Δ11, these are strong candidates to be exon 11 specific interactors.

In a complementary approach, we performed IP-MS on a cell line carrying an endogenous C-terminal triple TY-1 tag on BRCA1 (see Figure S2a-c for details on this cell line). In this experiment, we compared the BRCA1-interactome with and without DNA damage induction using ionizing radiation (IR) (Figure 1g, Supplemental Text and Supplemental Table 2). Again, we identified many of the known BRCA1 interactors, and most do not show an IR-dependent interaction, as has been observed before (Figure S2d) [22, 23, 36–39]. Interestingly, the strongest IR-dependent interactor, TOPBP1, also appeared in our other dataset as a significant interactor of FL BRCA1, but not BRCA1-Δ11 (Figure 1f,g). Given the previously described observations suggesting that TOPBP1 is involved in HR [19, 40, 41] and its previously described indirect interaction with BRCA1 via BRIP1[23] we decided to follow-up on this exon 11 mediated interaction.

We confirmed the interaction between exon 11 of BRCA1 and TOPBP1 by performing dedicated IPs in both directions using our virally complemented RPE1 *BRCA1*-KO cells upon IR treatment (Figure S3a and S3b). In addition, we validated the interaction in SUM149PT cells. This cell line, derived from a triple negative breast tumour, contains a frameshifting mutation in exon 11 which results in exon-skipping and expression of a *BRCA1-Δ11q* isoform [42]. Confirming our data in RPE1 cells, we did not observe an interaction between TOPBP1 and BRCA1-Δ11q upon IR in this more clinically relevant model. As a control, we virally expressed FL BRCA1 in these cells, which restored the interaction with TOPBP1 (Figure S3c). We also verified the interaction in mouse embryonic stem (mES) cells. For this, we used a previously described mES cell line carrying a conditional BRCA1 wildtype allele [29] and complemented these cells with human BRCA1 FL, Δ11 or 11q. Upon depletion of the wildtype mouse allele, the interaction with TOPBP1 was restored only in cells expressing FL BRCA1, and not the variants disrupting exon 11 (Figure S3d).

### BRCA1 contains a dual interaction mode for TOPBP1

Our IP-MS data showed that the interaction between BRCA1 and TOPBP1 is damage dependent. To confirm this, we performed a dedicated IP with and without IR in our RPE1 BRCA1-KO cells complemented with either FL BRCA1 or BRCA1-Δ11. Confirming our MS results, this IP shows IR-dependent binding of TOPBP1 to FL BRCA1 only, with no binding to BRCA1-Δ11 (Figure 2a). In the next sections where we have narrowed-down the interaction domains, all IPs were therefore performed after induction of DNA damage by IR. To rule out the possibility that the interaction between BRCA1 and TOPBP1 is driven by active resection and would therefore be reduced by BRCA1-Δ11 expression, we performed our IP experiments in a 53BP1 knock out setting as this has been shown to restore end-resection levels in BRCA1-Δ11 cells [35]. Also in this background, we see loss of TOPBP1 binding, indicating the interaction is not purely a consequence of active resection (Figure S3e). In total, our data indicate an IR-dependent interaction between TOPBP1 and BRCA1 mediated via exon 11, although biochemical experiments would be required to validate a direct interaction between TOPBP1 and BRCA1 exon 11.

**Figure 2.**
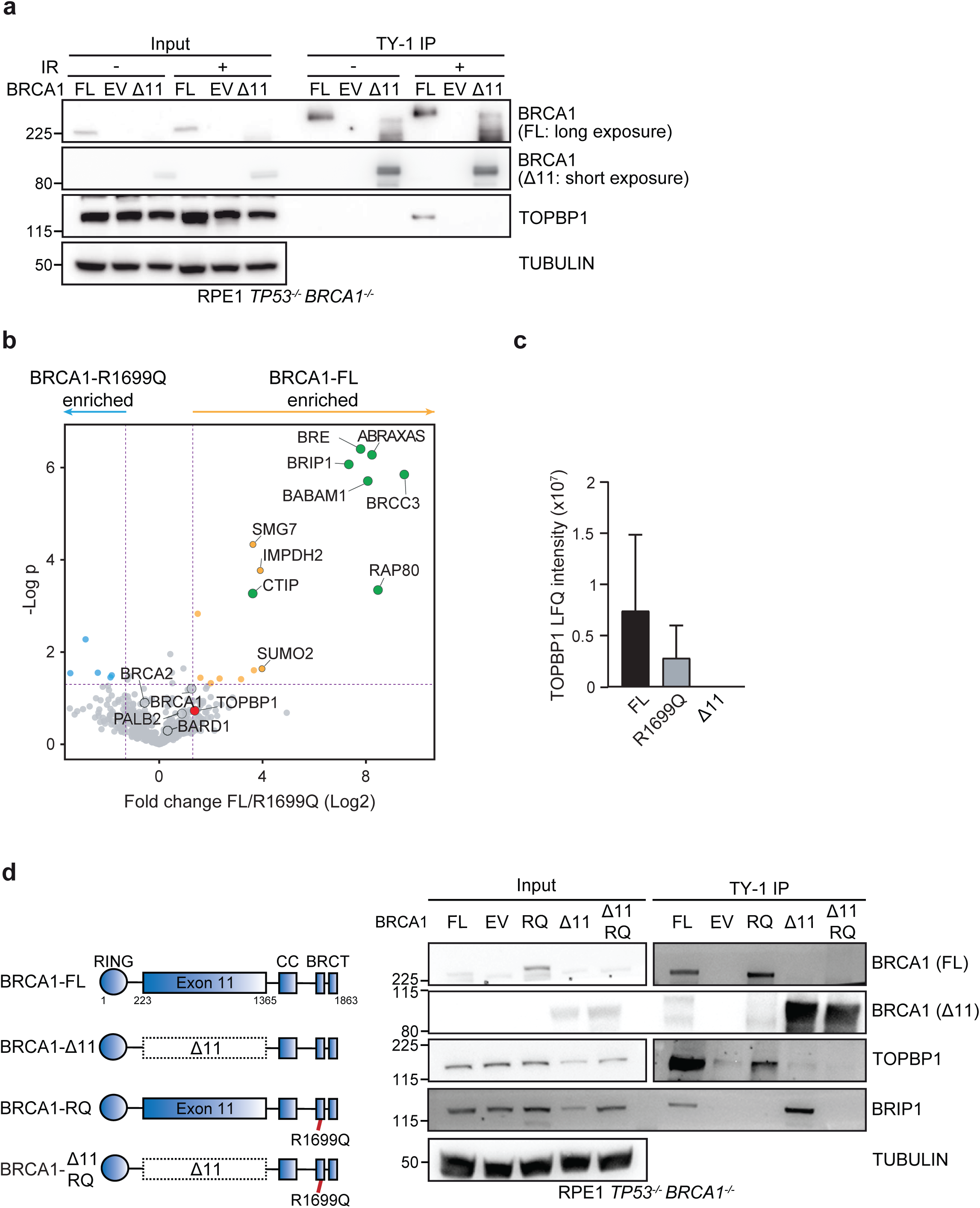
BRCA1 contains a dual interaction mode for TOPBP1. (A) Cell lysates from RPE1 hTERT *TP53^-/-^ BRCA1^-/-^* cells virally reconstituted with dox-inducible BRCA1-FL, BRCA1-Δ11 or EV control were subjected to TY-1 IP 1 hr post 5 Gy IR or noIR and analyzed by western blotting. (B) Volcano plot depicting statistical differences in binding proteins identified by MS proteomic analysis upon TY-1 IP of RPE1 hTERT *TP53^-/-^ BRCA1^-/-^* cells virally reconstituted with a dox-inducible BRCA1-FL or BRCA1-R1699Q 1 hr post 5 Gy IR (n=3). (C) Graph showing TOPBP1 LFQ intensity from the proteomic analysis of TY-1 IPs 1 hr upon 5 Gy IR of RPE1 hTERT cell lines *TP53^-/-^BRCA1^-/-^* virally reconstituted with BRCA1-FL, BRCA1-R1699Q and BRCA1-Δ11 (n=3, mean + SEM). (D) Schematic diagram of BRCA1-FL, BRCA1-Δ11, BRCA1 harboring a R1699Q mutation (RQ) or a double Δ11 and R1699Q mutation (Δ11, RQ) (left). Lysates of the indicated cell lines were harvested and IP’ed using a TY-1 antibody and analyzed by western blotting 1 hr post 5 Gy.

Previous research has shown that BRCA1 forms a complex with TOPBP1 via the bridging protein BRIP1 that binds the BRCA1 BRCT domains [23]. To better characterize the dual interaction modes between BRCA1 and TOPBP1, we generated cell lines expressing BRCA1 with a mutation in the BRCT (R1699Q) in the FL or Δ11 background. We performed IP-MS on the cells expressing FL BRCA1 R1699Q, which confirmed that this mutant disrupts the interaction with ABRAXAS, CtIP and BRIP1 (Figure 2b, Figure S3f, Supplemental Text and Supplemental Table 1). Quantification of the TOPBP1 peptides identified in the different MS datasets showed that TOPBP1 was most abundant in the

BRCA1 FL IP, followed by the BRCT mutant, whereas in the cells expressing BRCA1-Δ11 no peptides corresponding to TOPBP1 were identified at all (Figure 2c). These data indicate that the interaction between TOPBP1 and BRCA1 is most strongly regulated via exon 11. To confirm the proteomics data, we performed an IP on cells expressing the BRCT mutant either in the FL BRCA1 or Δ11 background. While disruption of either the BRCT or exon 11 reduced binding to TOPBP1, the effect was strongest in cells expressing BRCA1-Δ11 (Figure 2d). The double mutant did not show any binding of TOPBP1, indeed suggesting a dual binding mode of BRCA1 to TOPBP1. Importantly, the binding of BRCA1 to BRIP1 is functional in Δ11 cells, indicating that deletion of exon 11 does not lead to a disruption of the BRCT-mediated interactions (Figure 2d and Figure 1f and S1c).

### Characterization of the interaction between TOPBP1 and exon 11 of BRCA1

To further characterize the interaction between BRCA1 and TOPBP1, we generated deletion mutants in exon 11 by dividing the exon in four parts (Figure 3a). In an attempt to minimally affect protein folding, we used prediction software (RaptorX) to prevent disruption of alpha helix and beta sheet structures when creating the deletion constructs [43]. IP experiments with these deletion constructs showed reduced TOPBP1 binding to all of them, but only the deletion of the most C-terminal part of exon 11 completely disrupted binding (Figure 3a). We then further narrowed down the binding interface by creating 5 deletion constructs of this C-terminal part of exon 11 (Figure 3b). Deletion of the first or last 58 or 57 amino acids of this fragment did not affect binding, while all three deletion constructs of the middle part showed reduced binding indicating the binding to TOPBP1 is mediated via a minimal region comprising amino acids 1,134-1,307. Previous research has identified an SQ enriched region in BRCA1 (aminoacids: 1143-1525) [44] that coincides with the region that binds TOPBP1. Given the DNA damage-dependent nature of the interaction between BRCA1 and TOPBP1, we wondered whether ATM/ATR-dependent phosphorylation of the 4 SQ-sites within the TOPBP1-binding region was required for the binding. To study this, we generated a full length BRCA1 construct where all 4 Serines were mutated to Alanines and tested its interaction to TOPBP1. These mutations impaired binding to TOPBP1 compared to wildtype BRCA1 (Figure 3c), indicating that damage-dependent phosphorylation of exon 11 is required for its interaction with TOPBP1.

**Figure 3.**
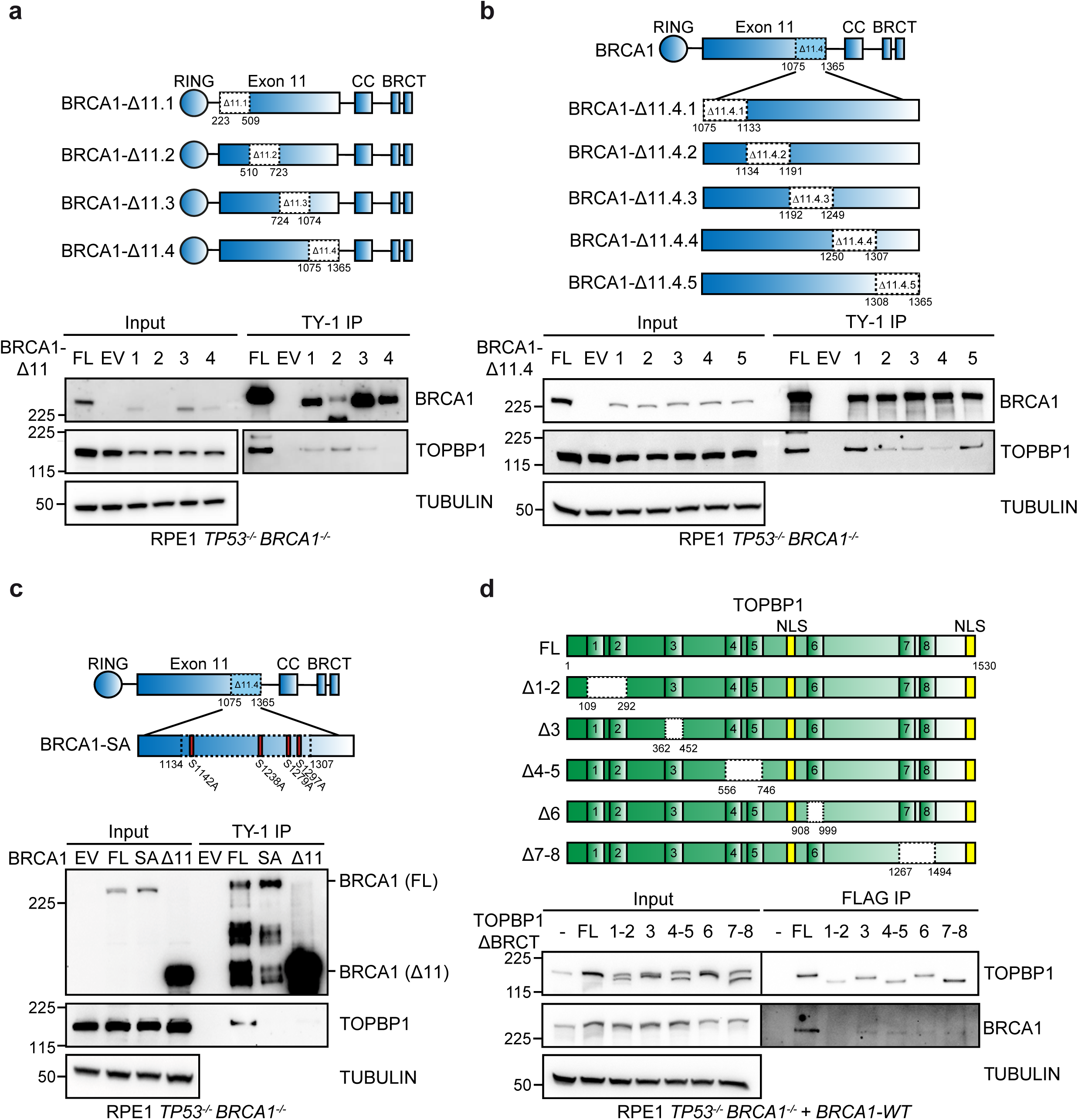
TOPBP1 BRCT1-2 bind to a C-terminal region of BRCA1 exon 11. (A) Schematic diagram of BRCA1 exon 11 deletion mutants used in this panel (top). Cell lysates from RPE1 hTERT *TP53^-/-^ BRCA1^-/-^* cells virally reconstituted with dox-inducible BRCA1-FL, the indicated exon 11 deletion mutants or EV control were subjected to TY-1 IP 1 hr post 5 Gy IR and analyzed by western blotting (bottom). (B) Schematic diagram of BRCA1 exon 11 C-terminal deletion mutants used in this panel (top). Lysates of the RPE1 hTERT *TP53^-/-^ BRCA1^-/-^* cells virally reconstituted with the indicated dox-inducible TY-1 tagged deletion mutants cells were subjected to TY-1 IP 1 hr post 5 Gy IR and analyzed by western blotting (bottom). (C) Schematic diagram of BRCA1-SQ sites in the C-terminus of exon 11 (top). IR-irradiated lysates (1 hr post 5 Gy IR) of the indicated cells virally reconstituted with dox-inducible BRCA1-FL or BRCA1-FL in which 4 SQ sites in the C-terminus of exon 11 were mutated to alanines (SA) were immunoprecipitated using a TY-1 antibody and analyzed by western blotting (bottom). (D) Schematic diagram of TOPBP1 single or tandem BRCT deletion mutants used in this study (top). IR-irradiated lysates (1 hr post 5 Gy IR) of RPE1 hTERT cell lines *TP53^-/-^ BRCA1^-/-^* virally reconstituted with dox-inducible BRCA1-FL and the indicated flag-tagged TOPBP1 deletion mutants were immunoprecipitated using a FLAG antibody and analyzed by western blotting (bottom).

*Vice versa*, we also investigated what domain of TOPBP1 directed the binding to BRCA1 exon 11. TOPBP1 consists of a series of eight BRCT domains, known for their interaction to phosphorylated proteins [45, 46]. The previously described interaction between BRIP1 and TOPBP1 occurs via BRCT 7 and 8 in TOPBP1 [21]. Additionally, a study using TOPBP1 point mutations has shown that BRCT1, 5 and 7 are involved in the interaction with BRCA1 [20]. To study which BRCT domains are involved in the binding to exon 11 of BRCA1, we created cell lines expressing different BRCT deletion constructs of TOPBP1 (Figure 3d). Immunoprecipitation of the different FLAG-tagged TOPBP1 deletion constructs showed that BRCA1 was able to bind to FL TOPBP1 and also the constructs lacking the BRCT regions 3 to 8 but not the construct in which the first two BRCTs were absent. This was confirmed by performing the IP in the reverse order by isolating BRCA1-TY-1 (Figure S3g). These results indicate that BRCA1 exon 11 binds TOPBP1 via a different region than BRIP1.

### The interaction between TOPBP1 and BRCA1 exon 11 is not involved in checkpoint activation

Given the well-described role of TOPBP1 in regulating ATR activity and cell cycle checkpoint activation [47–49], we investigated whether the interaction with BRCA1 is required for this activity. For this, we analyzed ATR-mediated phosphorylation events by western blot upon hydroxyurea (HU) treatment or IR in cells expressing BRCA1-Δ11 as a read-out for checkpoint activation upon replication stress or DSB induction, respectively. In both conditions, cells expressing FL BRCA1 or BRCA1-Δ11 showed similar ATR activation levels. These data indicate that the interaction between BRCA1 and TOPBP1 is not required for TOPBP1’s function in checkpoint activation (Figure S4).

### The BRCA1-TOPBP1 interaction mediates DNA end-resection during DNA repair

To study the biological relevance of the damage-induced interaction of BRCA1 with TOPBP1 in more detail, we set out to investigate whether the disturbed interaction with TOPBP1 is causative for the hypomorphic phenotypes of cells expressing BRCA1-Δ11. Depleting TOPBP1 using siRNAs had a small, non-significant effect on the recruitment of FL BRCA1 to damage sites, similar to the small decrease in recruitment of BRCA1-Δ11 (Figure S5a, Figure 1c). Moreover, TOPBP1 depletion did not further reduce recruitment of BRCA1-Δ11 (Figure S5a). Vice versa, our data show a clear reduction in TOPBP1 recruitment to sites of DNA damage in cells expressing BRCA1-Δ11 compared to full length BRCA1 (Figure S5b). In addition, the majority of the remaining TOPBP1 foci did not colocalize with BRCA1 (Figure S5b).

Cells expressing BRCA1-Δ11 show a strong reduction of pRPA foci formation, confirming the resection defect observed by others [13, 14, 35] (Figure 1d and 4a). siRNA-mediated depletion of TOPBP1 in cells expressing FL BRCA1 showed a similar reduction in pRPA foci as in cells expressing BRCA1-Δ11 (Figure 4a). Moreover, depletion of TOPBP1 did not further decrease pRPA foci formation in cells expressing BRCA1-Δ11, demonstrating an epistatic effect. Furthermore, ectopic expression of an siRNA-resistant TOPBP1 protein showed a rescue of the end-resection defects in TOPBP1-depleted FL BRCA1 cells. In contrast, this complementation did not show a recovery in BRCA1-Δ11 cells, indicating that the role of exon 11 is not just to regulate TOPBP1 protein levels but there is a mechanistic role for the interaction. Similar results were obtained when analyzing native BrdU staining as an independent read-out of end-resection (Figure S5c). To further validate a direct role of the TOPBP1-exon 11 interaction in DNA end-resection, we analysed pRPA foci upon re-expressing full length BRCA1 in which the 4 C-terminal SQ sites in exon 11 were mutated into SA. This mutant showed decreased end-resection with similar levels as BRCA1-Δ11 (Figure 4b). Altogether, these results show that TOPBP1 is involved in regulating end-resection via its interaction with exon 11 of BRCA1.

**Figure 4.**
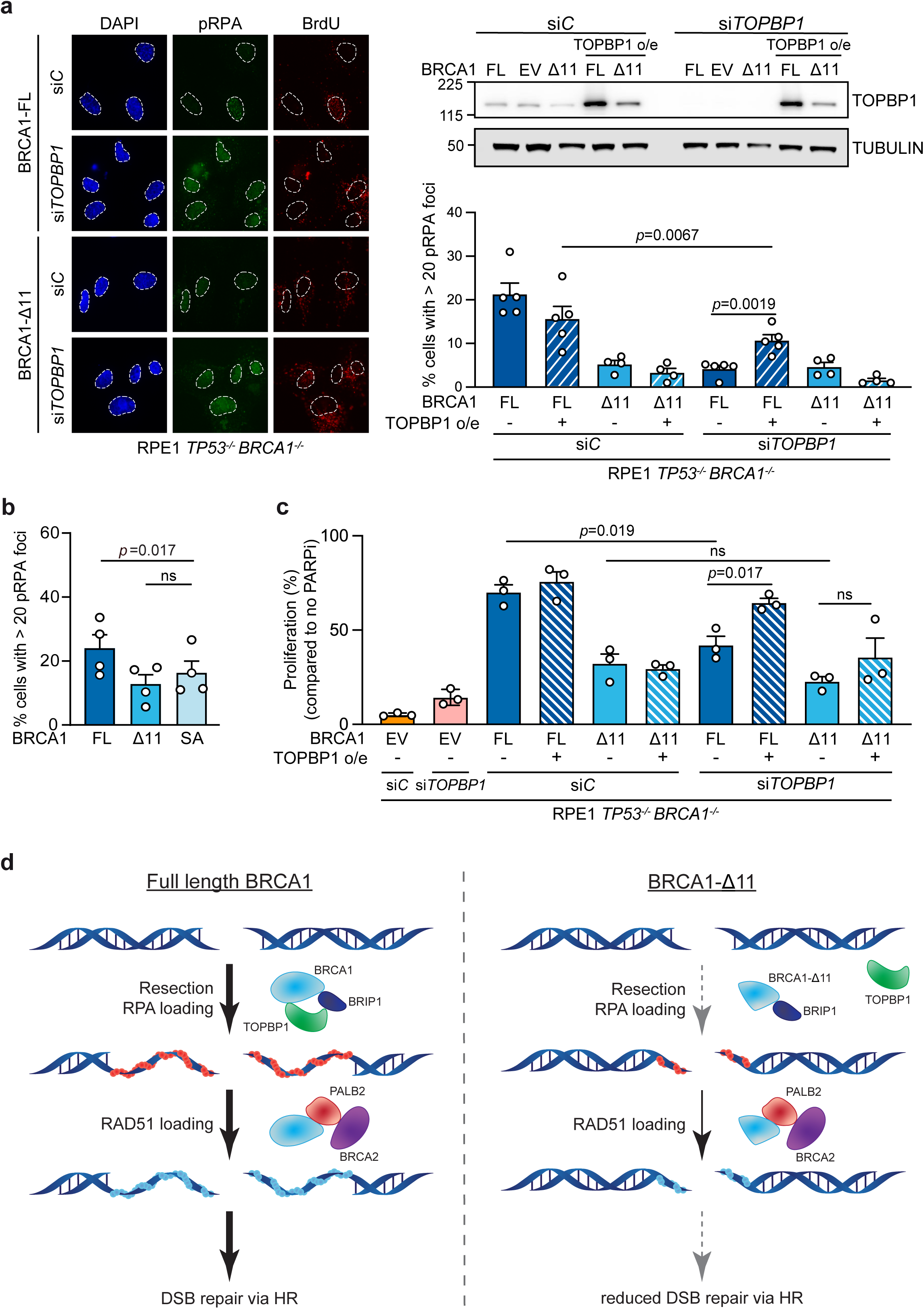
The BRCA1-TOPBP1 complex mediates DNA end-resection during DSB repair. (A) RPE1 hTERT *TP53^-/-^ BRCA1^-/-^* cell lines virally reconstituted with dox-inducible FL BRCA1, an EV control or BRCA1-Δ11 together with siRNA resistant FL TOPBP1 were transfected with a control (siC) or *TOPBP1*-targetting siRNA (siTOPBP1), followed by IF microscopy to analyze p-RPA and BrdU foci formation 4 hr post 10 Gy IR. Left panel shows representative microscopy images of p-RPA and BrdU foci. Upper right panel shows western blotting analyses of the cells mentioned above and lower right panel shows quantification of p-RPA foci (n=4/5, mean + SEM, paired t-test, p-values are shown). A minimum of 100 cells were counted per replicate. Quantification of BrdU signal is shown in figure S5c. (B) *TP53^-/-^ BRCA1^-/-^* virally reconstituted with dox-inducible TY-1 tagged FL BRCA1 wildtype or 4 x SA mutant or BRCA1-Δ11 were irradiated with 10 Gy and 4 hr post-IR analyzed by IF microscopy to detect pRPA foci formation (n=4, mean + SEM, paired t-test, p-values are shown). A minimum of 100 cells were counted per replicate. (C) Short-term proliferation assay of the cell lines described in Figure 4A treated with 1000 nM olaparib and dox. Data is normalized to cells treated with dox, but not olaparib (n=3, mean + SEM, paired t-test, p-values are shown). (D) Model of the TOPBP1-BRCA1 complex function during homologous recombination.

To test the impact of the interaction of BRCA1 with TOPBP1 on downstream HR, we studied PARP inhibitor (PARPi) sensitivity in our cell lines. Since TOPBP1 is an essential gene, we failed in generating clonal knock-out cell lines. Therefore, we depleted TOPBP1 using siRNAs and performed short-term PARPi sensitivity assays by measuring cell proliferation upon 7 days of olaparib treatment. western blotting confirmed that the used siRNAs against TOPBP1 resulted in TOPBP1 depletion for the duration of this experiment (Figure S5d). The depletion of TOPBP1 or the expression of BRCA1-Δ11 resulted in a similar sensitization of cells to PARPi and no additive effect was observed. The increased PARPi sensitivity by depletion of TOPBP1 via siRNAs could be rescued by overexpression of TOPBP1 in cells expressing FL BRCA1, but not BRCA1-Δ11 (Figure 4c). These results further align with a model in which the complex of BRCA1-TOPBP1 is important for efficient HR (Figure 4d).

## Discussion

Despite extensive studies on the function of BRCA1 over the last 30 years, there is still a substantial amount of information that remains unknown about its activities. A striking point about this protein is the fact that cancer-associated mutations occur throughout the gene and affect different protein domains and interactions, underlining its various roles in genome maintenance [50]. In this study we focused on the large exon 11 of BRCA1, which is often lost due to alternative splicing when mutations in exon 11 occur. This renders a protein lacking ∼60% of its full length while retaining all its major functionally characterized domains. Exon 11 contains two nuclear localization signals (NLS), however, previous research has shown that these are not required for nuclear localization of the protein [51, 52]. Indeed, in our model we observe normal recruitment of BRCA1-Δ11 to nuclear sites of DNA damage. Previous research on this small isoform has shown a hypomorphic function with a pronounced decrease in DNA end-resection upon DSB induction [9–11, 13, 14, 35, 53], which we also confirmed in our model.

To better understand the function of exon 11, we characterized its protein interactions and identified a novel DNA damage dependent interaction with TOPBP1. Our data show that the first two BRCT domains of TOPBP1 interact with a region containing 4 SQ sites in the C-terminal region encoded by BRCA1’s exon 11 (aa 1134-1307). Previous research has also shown that the first BRCT domain in TOPBP1 is involved in binding to BRCA1, confirming our data [20]. Mutating the serines of the 4 ATM/ATR SQ target sites into alanines prevented binding of TOPBP1 via exon 11, indicating that this interaction is phospho-dependent and hence could explain its DNA damage dependency. We were unsuccessful to further narrow-down the binding domain, suggesting there might be redundancy between these phospho-sites in regulating binding to TOPBP1. Previous research has shown that TOPBP1 and BRCA1 form a complex bridged by BRIP1 which binds both proteins [23]. This interaction involves BRCT 7 and 8 of TOPBP1 [21, 54], indicating that TOPBP1 contains two binding modules for BRCA1. Our data confirm this dual mode of interaction since deletion of exon 11 does not fully prevent complex formation between BRCA1 and TOPBP1, although the interaction via exon 11 seems to be the most dominant form in our model.

We show that the interaction between TOPBP1 and the exon 11-encoded region is required for TOPBP1 recruitment to DSBs and mediates BRCA1’s role in end-resection during DSB repair. This confirms previous research that showed that TOPBP1 is involved in DSB repair and HR by driving end-resection [19, 20, 55], although others have claimed a resection independent role of TOPBP1 in HR [40]. TOPBP1 also plays an important role in DSB repair during mitosis [56–58]. This function, however, seems independent of its function with BRCA1 since BRCA1 is not recruited to sites of damage during mitosis [59, 60]. Besides a role in HR, TOPBP1 is most well-known for its role in checkpoint and ATR activation [16, 47, 49]. Our results suggest that this function does not require an interaction with BRCA1, as BRCA1-Δ11 cells do not show checkpoint defects.

In total, our data indicate that the role of exon 11 of BRCA1 in end-resection is mediated via an interaction with TOPBP1. These data provide further understanding of TOPBP1’s role during HR, although the mechanistic role of the BRCA1-TOPBP1 complex during end-resection will need to be explored in more detail in the future.

## Acknowledgements

This work was financially supported by Oncode Institute funds for SMN, a Dutch Research Council (NWO) VIDI grant (VI.Vidi.192.039) to SMN, a Dutch Cancer Foundation (KWF) young investigator grant (11367 / 2017-2) to RGP, and an NWO Medium Investment Grant (91116004) to PAV.

## Author Contributions

The study was designed by SMN with input from MSMA. All experiments were performed by MSMA, with help of RK, AS, and VG. RM provided mES cell models. Mass spectrometry runs were conducted by AHR, supervised by PAV. Proteomic analyses were performed by DS, supervised by RGP. The manuscript was written by MSMA and SMN, with input from all co-authors.

## Data availability

All raw data will be deposited on Mendeley data and proteomic datasets will be uploaded to ProteomeXchange upon acceptance.

Figure S1. **BRCA1 exon 11 mediates HR and interacts directly to TOPBP1 (related to figure 1)** Representative microscopy images of p-RPA (A) or RAD51 foci (B) quantified in Figure 1B and 1C, respectively. (C) Volcano plots depicting statistical differences in binding proteins identified by MS proteomic analysis upon TY-1 IP of RPE1 hTERT TP53^-/-^ BRCA1^-/-^ cells virally reconstituted with dox-inducible BRCA1-FL or BRCA1-Δ11 1 hr post 5 Gy IR compared to EV 1 hr post 5 Gy IR (n=3).

Figure S2. **Generation of an endogenous BRCA1-TY-1 cell line and analyses of its interactors by Mass Spectrometry (related to figure 1)** (A) Schematic representation of the approach to knock-in a C-terminal triple TY-1-P2A-neomycin tag into the human 3’ *BRCA1* locus (top) in RPE1 *TP53^-/-^* cells and Sanger sequencing results of the region of the integration (bottom). (B) Lysates of RPE1 hTERT cell lines either *TP53^-/-^* or *TP53^-/-^ BRCA1-TY-1* endogenously tagged analyzed by western blotting to visualize BRCA1 protein levels. (C) Representative images of RPE1 hTERT *TP53^-/-^* endogenously tagged BRCA1-TY-1 cells IR-treated (10 Gy) and analyzed 4 hr post IR to detect TY-1 and RAD51 foci. (D) Volcano plots depicting statistical differences in binding proteins identified by MS proteomic analysis upon TY-1 IP of RPE1 hTERT *TP53^-/-^ BRCA1-TY-1* compared to RPE1 hTERT *TP53^-/-^* cells not irradiated (left) or 1 hr post 5 Gy IR (right).

Figure S3. **Validation of the interaction between TOPBP1 and BRCA1 exon 11 in different cell systems (related to figure 1, 2 and 3)** (A) Cell lysates harvested 1 hr post 5 Gy IR from RPE1 hTERT *TP53^-/-^ BRCA1^-/-^* cells virally reconstituted with dox-inducible TY-1 tagged BRCA1-FL, BRCA1-Δ11 or an EV control were subjected to TY-1 IP and analyzed by western blotting. (B) Cell lysates harvested 1 hr post 5 Gy IR from RPE1 hTERT *TP53^-/-^ BRCA1^-/-^* cells virally reconstituted with dox-inducible TY-1 tagged BRCA1-FL, BRCA1-Δ11 or an EV control were subjected to endogenous TOPBP1 IP and analyzed by western blotting. (C) SUM149PT cells virally reconstituted with dox-inducible TY-1 tagged BRCA1-FL or EV were subjected to TOPBP1 IP 1 hr post 5 Gy IR and analyzed by western blotting. (D) Lysates of BRCA1-deficient mES cell lines reconstituted with human BRCA1-FL, -Δ11 or −11q were subjected to TOPBP1 IP 1 hr post 5 Gy IR and analyzed by western blotting. (E) Cell lysates harvested 1 hr post 5 Gy IR from RPE1 hTERT *TP53^-/-^ BRCA1^-/-^* or *TP53^-/-^ BRCA1^-/-^ 53BP1^-/-^* cells virally reconstituted with dox-inducible TY-1 tagged BRCA1-FL, BRCA1-Δ11 or an EV control were subjected to TY-1 IP and analyzed by western blotting. (F) Volcano plots depicting statistical differences in binding proteins identified by MS proteomic analysis upon TY-1 IP of RPE1 hTERT *TP53^-/-^ BRCA1^-/-^* cells virally reconstituted with dox-inducible BRCA1-R1699Q 1 hr post 5 Gy IR compared to EV 1 hr post 5 Gy IR (n=3). (G) Cell lysates harvested 1 hr post 5 Gy IR of RPE1 hTERT TP53^-/-^ BRCA1^-/-^ cells virally reconstituted with dox-inducible BRCA1-FL and the indicated FLAG-tagged TOPBP1 deletion mutants were subjected to TY-1 IP and analyzed by western blotting. Please note that the TOPBP1 staining also visualizes endogenous TOPBP1. For the IP samples, all lanes show a combination of endogenous and exogenous protein variants, indicating normal binding of the mutant, except the deletion mutant of BRCT1-2, which does not appear in the IP sample.

Figure S4. **The interaction between BRCA1 exon 11 and TOPBP1 is not required for checkpoint activation (related to figure 4)** Cell lysates from RPE1 hTERT *TP53^-/-^ BRCA1^-/-^* cells virally reconstituted with dox-inducible BRCA1- FL, BRCA1- Δ11 or EV control and transfected with a control (siC) or *TOPBP1*-targetting siRNA (siTOPBP1) harvested 6 hr post 20 mM HU treatment (top) or 6 hr post 5 Gy IR (bottom) were analyzed by western blotting using the indicated antibodies.

Figure S5. **The interaction between BRCA1 and TOPBP1 stimulates DNA end-resection (related to figure 4)** (A) Cell lysates harvested 1 hr post 5 Gy IR from RPE1 hTERT *TP53^-/-^ BRCA1^-/-^* cells virally reconstituted with dox-inducible BRCA1-FL, Δ11 or EV control and transfected with a control (siC) or *TOPBP1*-targetting siRNA (siTOPBP1) were analyzed by western blotting to check correct siRNA depletion (left). Representative microscopy images (middle) and quantification (right) of BRCA1-TY-1 foci formation 1 hr post 5 Gy IR of the same cells (n=3, mean + SEM). A minimum of 100 cells were counted per replicate. (B) Representative microscopy images of TOPBP1 and BRCA1-TY-1 foci from RPE1 hTERT *TP53^-/-^ BRCA1^-/-^* cell lines virally reconstituted with dox-inducible FL BRCA1, an EV control or BRCA1-Δ11 1 hr post 5 Gy. Top right graph shows quantification of TOPBP1 foci (n=3, mean + SEM, paired t-test, p-values are shown). A minimum of 100 cells were counted per replicate. Bottom right graph shows TOPBP1+BRCA1 colocalization (n=3, mean + SEM, paired t-test, p-values are shown). A minimum of 25 cells were counted per replicate. (C) Quantification of BrdU foci from cells described in Figure 4A (n=4/5, mean + SEM, paired t-test, p-values are shown). A minimum of 100 cells were counted per replicate. (D) Cell lysates from RPE1 hTERT *TP53^-/-^ BRCA1^-/-^* cells virally reconstituted with TY-1 tagged BRCA1-FL transfected with a control (siC) or *TOPBP1*-targetting siRNA (siTOPBP1) were analyzed by western blotting (related to Figure 4B).

