## Supplementary figures and images for "DNA end-resection is stimulated by an interaction between BRCA1 exon 11 and TOPBP1"

### Supplemental figures

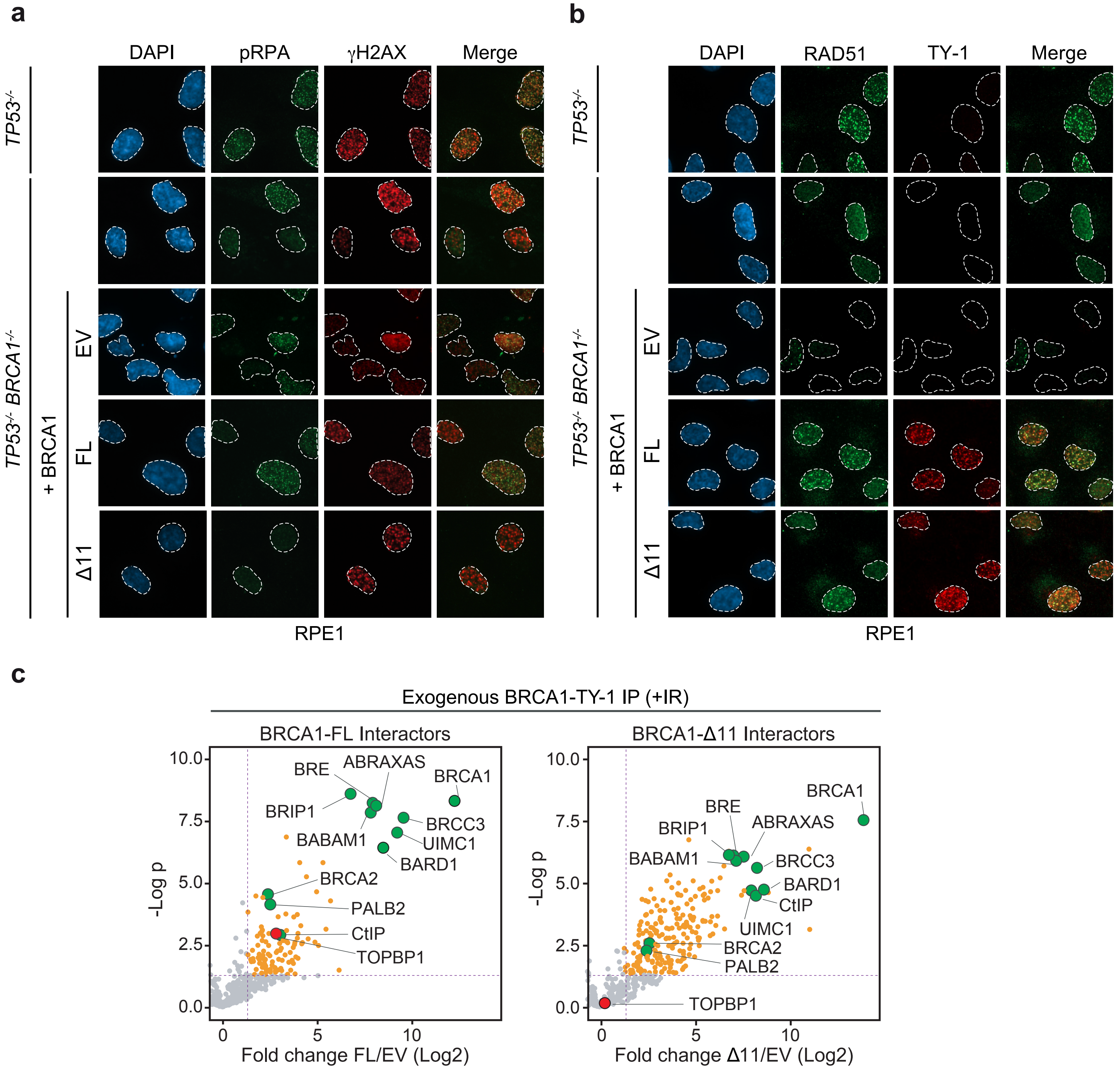

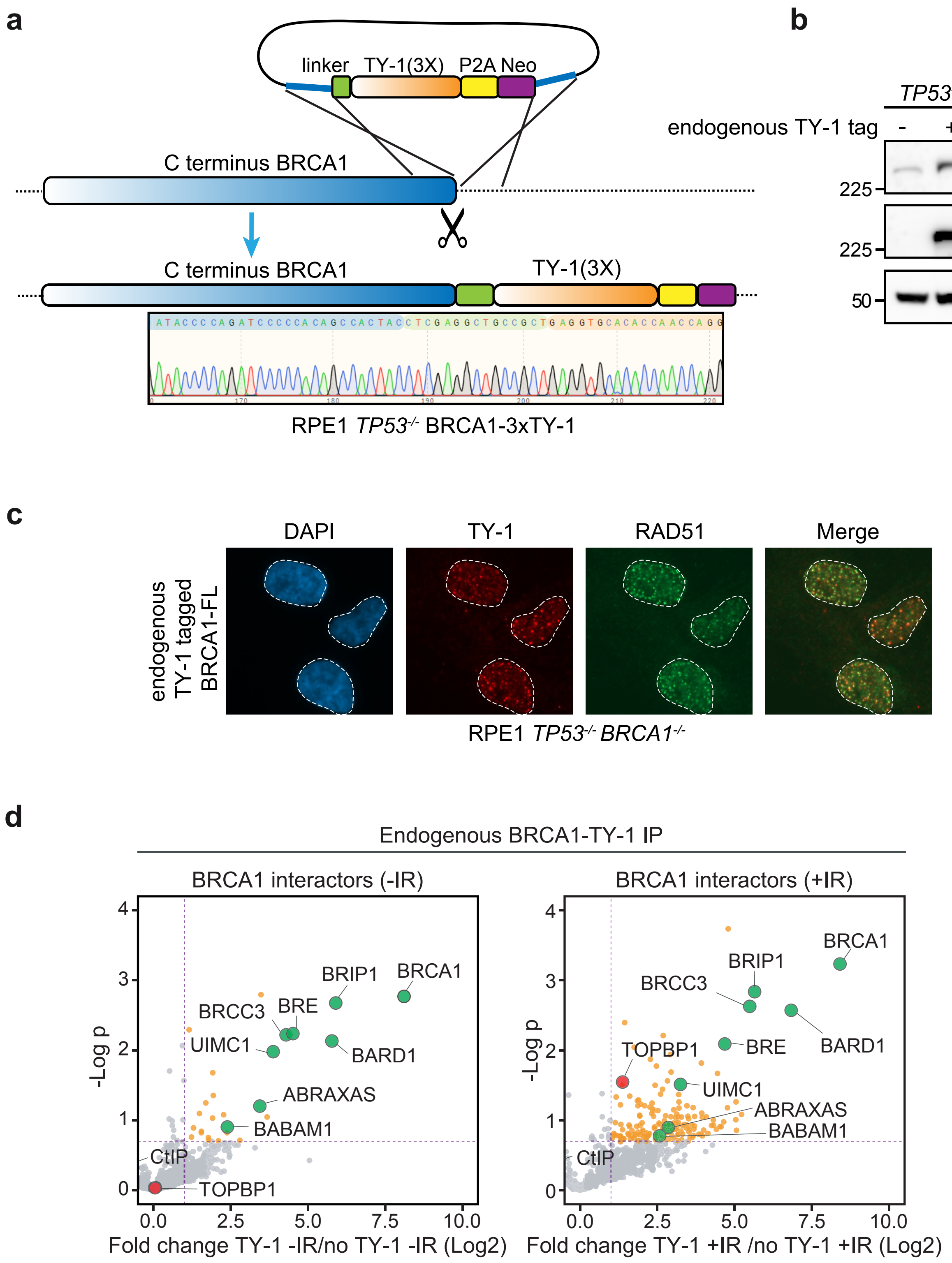

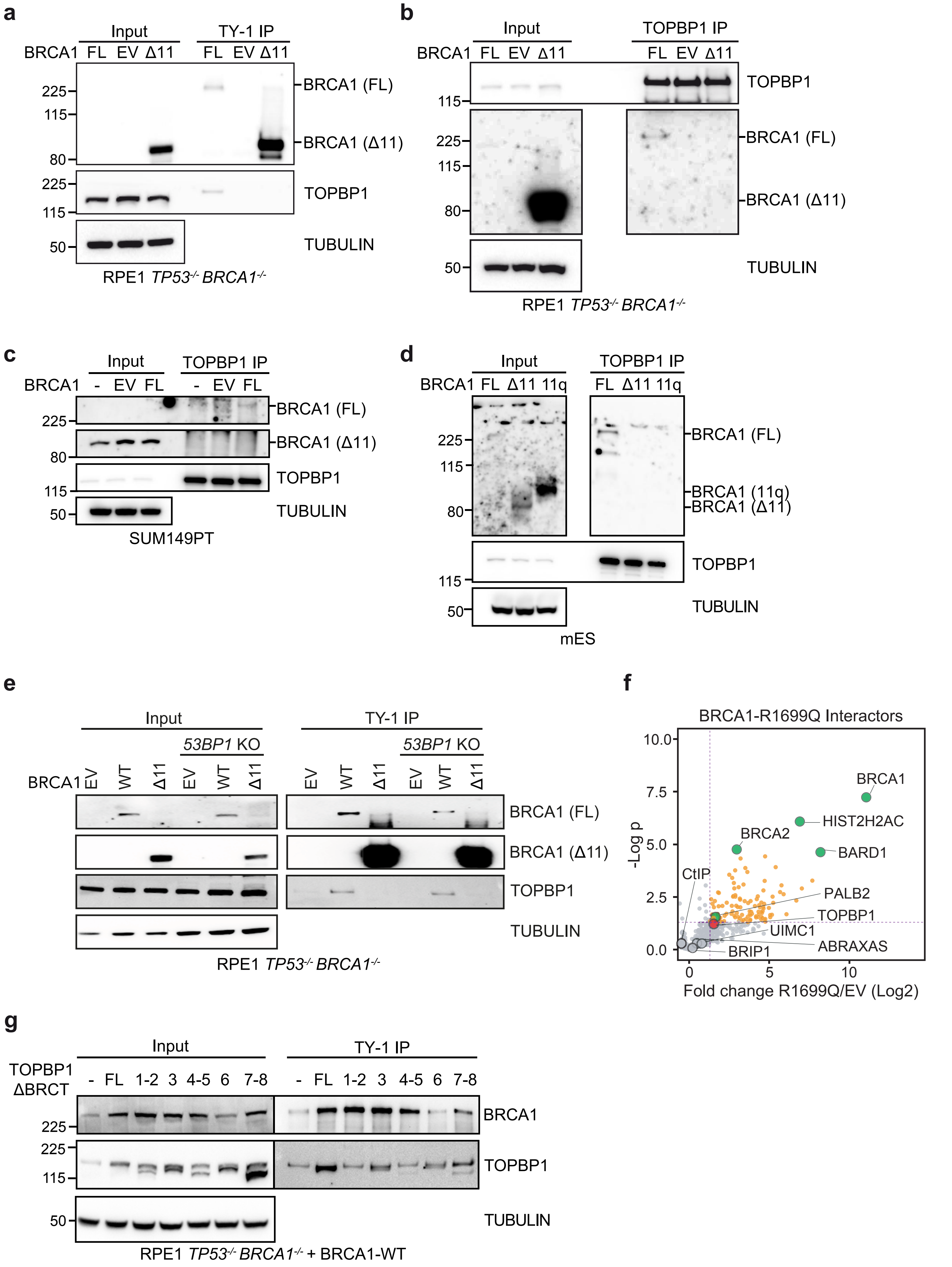

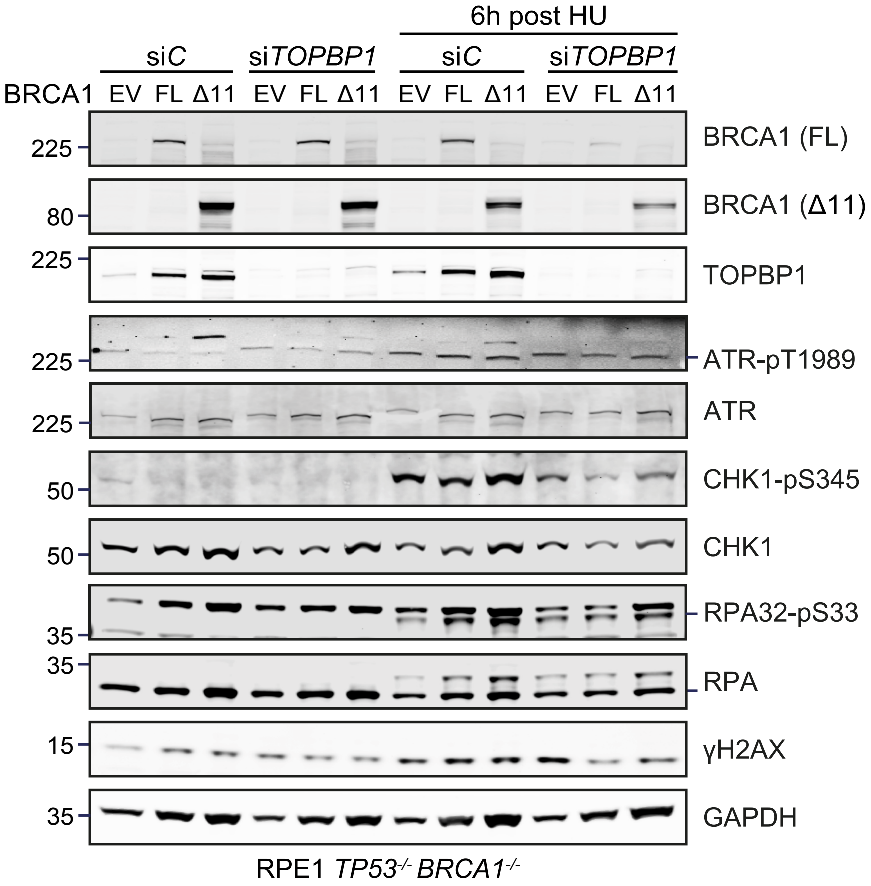

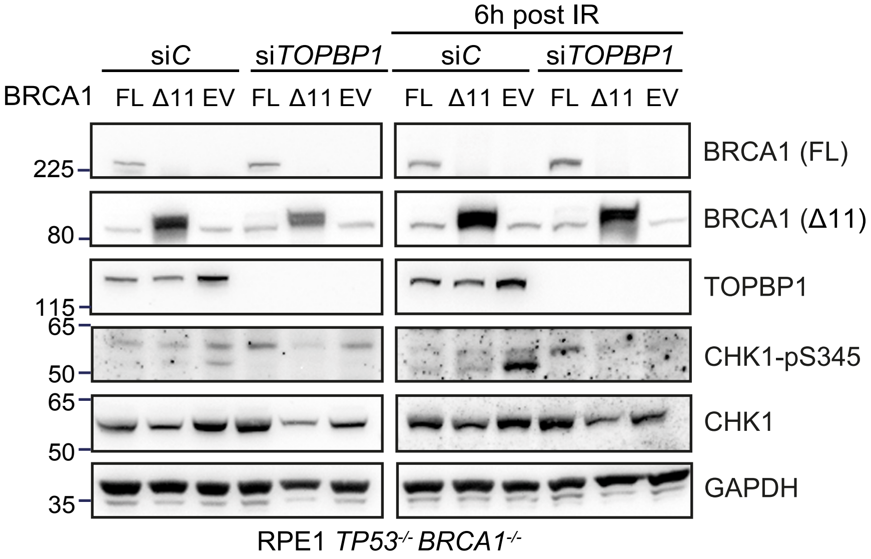

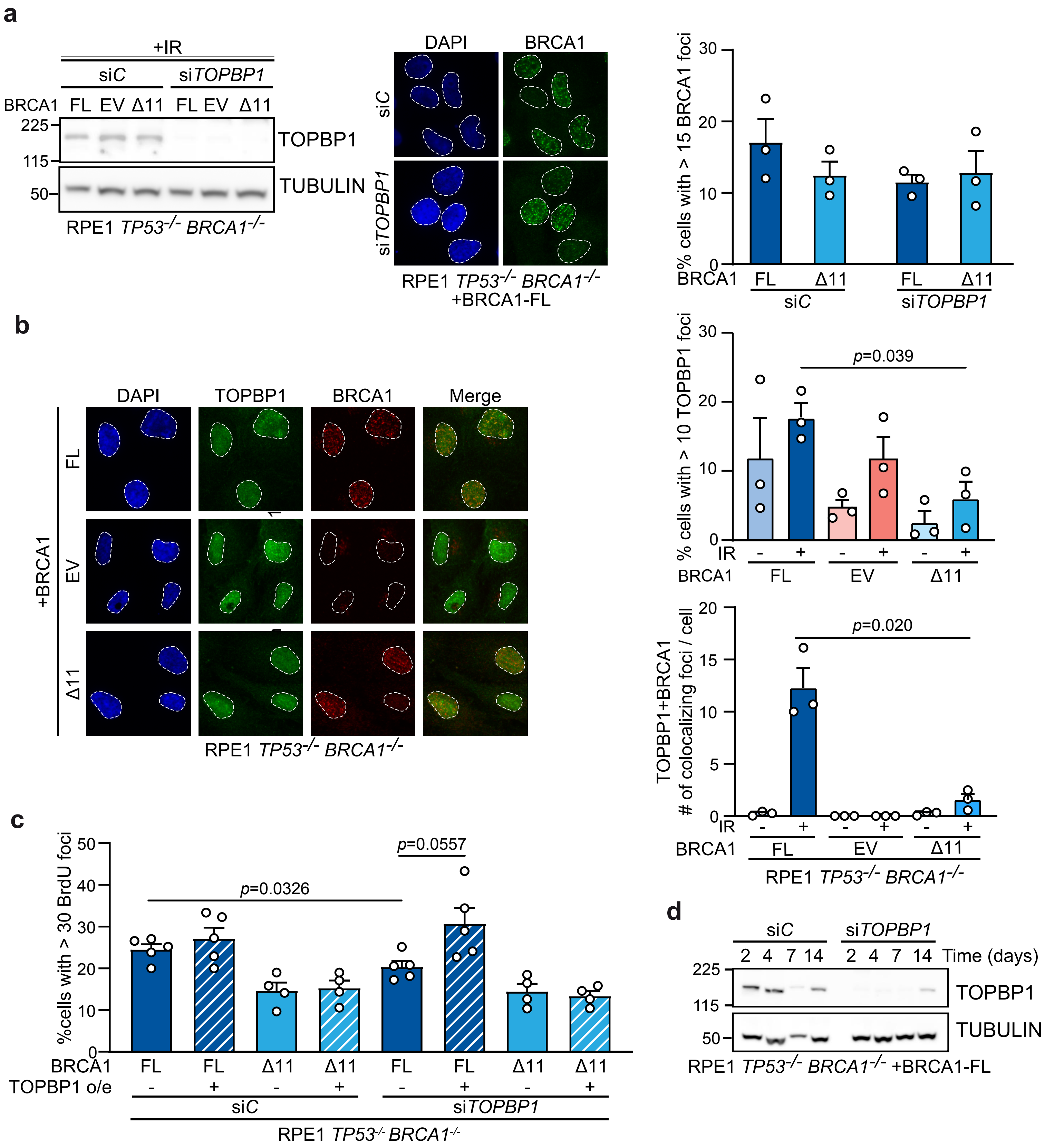
