## Supplemental text for "DNA end-resection is stimulated by an interaction between BRCA1 exon 11 and TOPBP1"

**Explanatory note on the proteomics result tables (supplemental table 1 and 2)**

Reading the tables from left to right:

- the first 3 columns list the gene names and protein names of all proteins identified in this MS screening.
- Next, columns are provided with a simple YES/NO to indicate whether a protein is considered a BRCA1 interactor (BRCA1 WT, BRCA1-Δ11 or BRCA1-R1699Q for Supplementary table 1; IR or no IR for Supplementary table 2). Proteins are considered interactors if they pass a Student's T-test ($p<0,05$) and show a log2 fold change > 1 over empty vector (S1) or untagged cells (S2) (i.e, fold change > 2).
- Subsequently, results from statistical analysis comparing two groups as indicated in title using PERSEUS software are shown:
    - Student's T-test: The PERSEUS software gives a "+" for proteins that are statistically significant ($p<0,05$).
    - Log Student's T-test p-value: This is the p-value given by PERSEUS software in -Log10 format.
    - Student's T-test Difference. This represent the Log2 fold change between two groups.
- The miscellaneous columns provide details on the peptides identified per protein ((total, razor and unique peptides), sequence coverage of the protein (total, razor and unique peptides), molecular weight of the protein, Q value (the ratio of reverse to forward protein groups for FDR control), Score (derived from peptide posterior error probabilities), intensity (Summed up eXtracted Ion Current (XIC) of all isotopic clusters associated with the identified protein), iBAQ peptides, MS/MS counts).
- The LFQ values refer to the Label-Free Quantification value calculated using the MaxLFQ algorithm.
